# Historical demography and species distribution models shed light on past speciation in primates of northeast India

**DOI:** 10.1101/2023.02.25.530015

**Authors:** Mihir Trivedi, Kunal Arekar, Govindhaswamy Umapathy

## Abstract

**Objectives:** To ascertain the effect of historical demography and past climate change as the drivers of diversity in northeast India.

**Materials and methods:** We took the variant called whole genome files of nine species present in the northeast India from Primate genome sequencing consortium work and assessed each species historic effective population size by using Multiple Sequentially Markovian Coalescent (MSMC) tool. We also constructed species distribution models on past (Pliocene and Pleistocene) and present climate with Maxent, by utilizing publicly available distribution data for each species.

**Results:** We got the effective population sizes for 10 million years ago at most, though we considered the data only till 3.3 million years. All species showed rise and decline at various time periods. The species distribution models showed disparate distribution at all three time points with a genera-wise pattern emerging.

**Discussion:** We found that the evolutionary trajectories of all the four genera into consideration, *Macaca, Trachypithecus, Hoolock* and *Nycticebus* are different from each other. Species in *Macaca* looks to be evolved in northeast as well as come from southeast Asia. Some species of *Trachypithecus* seems to radiated in the northeast India. Similarly, *Hoolock* has evolved in the region and *Nycticebus* is predicted have arrived from Indochina in the region. Hence, this study provides unique insights to the evolutionary dynamics for primate species in the northeast India.

## Introduction

Recent global climate change has brought attention to the impact of climate on the living world, with the major question being focussed on its effects on individual species, but before that we need to understand the effect of climate change on biodiversity in general. Climate change can have a drastic effect on adaptation of a population and in the long term can also give rise to speciation (Hoffmann & Sgró, 2011; Hua & Wiens, 2013; Parmesan, 2006). This correlation between climate and speciation has garnered more attention in recent years after the awareness of human mediated climate change has permeated both scientific and popular discourse (Jaureguiberry et al., 2022; Lenoir et al., 2017; Midgley et al., 2002). Due to anthropogenic factors influencing climates, many species have been documented to move towards favourable habitats (Evans et al., 2020; Neumann et al., 2022; Poessel et al., 2022). It has been hypothesized that any influence of climate on rates and patterns of speciation can be one of the important causes of present patterns of biodiversity, particularly the high richness of tropical regions (Mittelbach et al., 2007). Even a small magnitude of change in climate, if persistent for long, can influence natural histories of species and their spatial extent in a particular area (Parmesan & Yohe, 2003).

Such presence of related species in disparate regions can be explained by two alternative processes: vicariance and dispersal. In vicariance events, the speciation is caused by formation of physical barriers and therefore one can give information about the other. But in dispersal scenario, geographical barriers predate any speciation events. The widespread acceptance of the theory of plate tectonics brought vicariance as a possible mechanism for segregation of populations and speciation on the forefront (Briggs, 1987; Trewick, 2017).

Another kind of dispersal is when a species enlarges its distribution, because of conducive climatic or geographical changes. These can be termed as geo dispersal events (Lieberman & Eldredge, 1996). As opposite to vicariance, various regions get more connected here. This can be very well true for Indian biogeography, as Indian plate after collision form Asian plate gave opportunity to species in southeast Asia to spread towards the subcontinent, with present day northeast India as their doorway. Species’ range shifts have occurred throughout the Earth’s history. For instance, historical glacial cycles have caused range shift in many species (Beyer & Manica, 2020; Davis & Shaw, 2001; Jackson & Overpeck, 2000). Therefore, temperature plays a major role in deciding species ranges. Recent research has shown that this is also true for aquatic ecosystems.

In drastic ecological changes like temperature variations and glacial periods, a population can save itself from extinction by moving in a more suitable area, or employing phenotypic plasticity or undergoing evolutionary adaptations which may result in speciation (Williams et al., 2008). Therefore, dramatic climate change can give rise to a variety of selection pressure on a population, or on the species. For example, thermal stress produces directional selection on species which cannot regulate their body temperatures like porcelain crabs and lizards (Huey et al., 2009; Stillman, 2002). Similarly, some species have moved towards suitable habitats in response to climate change (Crispo et al., 2010). Range boundaries are the locations where most biotic and abiotic interactions happen. These interactions can give rise to strong selection pressures, and also population bottlenecks. The feedback between ecology and evolution is strongest at range boundaries, where selection is assumed to be strongest and where population bottlenecks are common (Hill et al., 2011). Inter- and intra-specific interactions in a community can be affected by resource availability, propagule availability, competition, predation, symbioses, and facilitation. These factors in the long term can be the reasons for speciation of populations. Climate change can play a predominant role in the aforementioned processes by disturbing established interactions or by giving rise to new interactions (Lavergne et al., 2010).

Today we have a better understanding of biodiversity due to satellite cartography and GIS systems. This privilege of technology has also enabled us to model the distribution of a species. Species distribution models predict the distribution of a species using presence or absence data and the values of environmental variables at these sites (Hao et al., 2019). These models can be extrapolated in space and time to infer distribution at a particular place and at a particular time. The climatic conditions in which a species occurs are used to construct a model of the species’ ecological niche that is then projected into geographical space (Guisan & Thuiller, 2005; Teixeira et al., 2021).

Here, we concentrated mainly on two epochs, which are the most recent ones: Pleistocene and Pliocene. Pliocene had a warmer climate than present but colder to the preceding Miocene. This resulted in a broader distribution of plant and animal species due to survivable warm conditions in norther latitudes, especially near the Arctic in the Northern hemisphere. In Pliocene the highest temperatures were reached between 3 and 4 million years ago (Haywood et al., 2009). This change in vegetation pattern became a key factor in the evolution of various animals, like ungulates and their predators (McKee, 2001; Salzmann et al., 2008). It has been showed that this makes Pliocene a suitable model system for future climate prediction scenarios (Tierney et al., 2020). In Pleistocene, the warmer world of Pliocene cooled down and gave rise to glacial/interglacial cycles, which continued till the Last Interglacial (LIG) ending at 11,700 years ago. Pleistocene ecosystems were very close to the present ecosystems. This climate was conducive of various plants and animals with high nutritious biomass which support animals such as woolly mammoths and great teratorn birds. Many of the animals which thrived in Pleistocene are still surviving to this day. But along with a favourable climate in interglacial, the conditions also became severe during the glacial, which resulted in local exterminations of the species, as well as gave selective pressures for speciation (Hewitt, 2000; Smith et al., 2022).

The Northeast India is at the confluence of mainland south Asia and Southeast Asia. This makes it a unique region to study the species’ range dynamics and speciation. With fourteen documented primate species in the region, it is an extraordinarily diverse area. We investigated 9 primate species from the region to elucidate the reasons for such diversity. Our approach included two separate dimensions: 1. to understand the demographic history of individual species; 2. to model their historic distribution by extrapolating current geographical locations. Demographic history will provide the fluctuations in effective population size which has been implicated in speciation events (Chen et al., 2020; Harvey et al., 2019). Similarly, climatic changes alter the selective pressure on a population, which in turn can be an impetus for diversification in a population (Barnosky & Kraatz, 2007).

These complementary approaches, along with published divergence details, can provide us a holistic understanding of how various species either evolved or arrived from different locations in present day northeast India. We investigate that how the species’ distribution in Pliocene and Pleistocene correlates with the effective population size and how this relation can provide insights upon the drivers of primate diversity in the northeast India.

## Materials and Methods

### Historical demography

We took the samples from nine species of primates, with individuals varying from 1 to 11, as shown in Table 4.1. The VCF files were taken from the publication Kuderna et al. (Unpublished data, in review). If the whole assembled genome of the same species was not available, then variants were called by referencing it to another related species. Callable ‘bed’ and variant ‘vcf’ were downloaded from the database and both were split separately as per chromosome or contig structures. Input files were generated by combining these files with generate_multihetsep.py command provided in MSMC tools. MSMC2 was run with these input files. The output was plotted with generation times and mutation rates given in Table 4.2.

### Palaeo-distribution modelling

#### Data collection

The occurrence data for the nine species was collected from the GBIF Global Biodiversity Information Facility) database (See references for DOI of each dataset). For each species, we plotted these occurrence records on the map and included only those records which fell within the known distribution zones of the respective species as per IUCN (iucnredlist.org). In some instances, the GBIF dataset had multiple points for the same location. Therefore, in order to obtain a single occurrence record per location, we used the spThin package in R v4.0 (Aiello-Lammens et al., 2015).

Further, we downloaded the 19 bioclimatic variables from paleoclim.org (Brown et al., 2018). We also downloaded layers for three different climatic periods – current (1979-2013), Pleistocene MIS19, ca. 787 ka (thousand years ago) (Brown et al., 2018) and Pliocene M2, ca. 3.3 Ma (million year ago) (Dolan et al., 2015). Both the past layers, i.e., the Pleistocene and Pliocene, contained only 14 bioclimatic layers – bio1, bio4, bio8, bio9, bio10, bio11, bio12, bio13, bio14, bio15, bio16, bio17, bio18, bio19, because the remaining layers could not be created for these time periods, as mentioned in paleoclim.org. Therefore, for the current time period as well, we used the same 14 layers and removed the rest. Spatial resolution of all the layers was 2.5 arcmins. Using ArcMap v10.2.2, all the bioclimatic variables were clipped for the region from 67.12 °E to 124.12 °E and from 12.6 °S to 39.35 °N. The clipped layers were exported into ASCII format for further use. These layers were then checked for multicollinearity using the correlation option in ENMtools v1.3 (Warren et al., 2010).

#### Variable selection

For each of the 9 species, a test run was performed using Maxent v3.4.3 (Phillips et al., 2006). Here, we used layers from only the current time period, all the 14 bioclimatic variables were used for this test run. Default values were used for the run except for the following modifications; feature type=Auto, RM (Regularisation multiplier) =1, maximum iterations=5000, replicates=10 and replicate type =subsample. After the analysis, only those variables were chosen whose percent contribution was greater than 1%. From these chosen variables, only the variables with the correlation (R) ≤ 0.75 were selected for the final analysis.

#### Model selection

The complexity of Maxent models is controlled by type of feature class (FC) and the value of regularisation multiplier (RM) (Morales et al., 2017; Radosavljevic & Anderson, 2014) which in turn controls the quality of the Maxent output (Shcheglovitova & Anderson, 2013). Therefore, to find the best combination of FCs and RM value, we performed this model selection. Maxent v3.4.3 has the following FCs - linear (L), quadratic (Q), product (P), hinge (H), threshold (T), and Auto (Philips et al., 2017), whereas the default RM value is 1. We used 11 sets of FCs – H, L, LQ, LQH, LQP, LQPT, LQPTH, Q, QPT, QPTH, and T. These 11 sets resulted from either a single FC or a combination of multiple FCs. Then for each of these 11 FCs, we used four different RM values – 0.5, 1, 2, and 5. In total, for each of the 9 species, we tested 44 different models (model here is a combination of FC and RM value). The best model was selected (Table 4.3) by performing the model selection analysis in ENMTool v1.3 using the AICc criteria (Galante et al., 2018).

#### Maxent analysis

For all the 9 species of primates, the final analysis was performed in Maxent v3.4.3 using the following modifications; random test percentage was set to 25%, maximum number of background points was set to 10000, the replicates were set to 10 and the replicate type was changed to sub-sample. 5000 iterations were performed, Jackknife test was performed to estimate the contribution of each environmental variable. The FC type and RM values were different for each species and were selected based on the model selection analysis (Table 4.3). The output format was chosen as Cloglog. AUC values were examined to check for the predictive ability of the model built.

## Results

### MSMC2 estimates

For all the species, we could get the estimates of effective population size till at least the beginning of Pleistocene. Some species data was also showing the estimates going it 10 million years ago, though we did not take them into consideration as climate data for such ancient time period is not available. All the four species of *Macaca* are very varied, with different patterns rise and decline in population sizes. Among the langurs, *Trachypithecus pileatus* shows a rise at about 3.3 million years point and then declines. *T.geei* data shows a rise at mid-Pleistocene and declines after that. The pattern is similar in *T. phayrei. Hoolock hoolock* had equally large population comparative to recent times, with highest point at about 800,000 years ago and has declined since. *Nycticebus bengalensis* also has the highest number of individuals at mid Pleistocene and since has declined as Holocene has come along.

### Maxent models

The bioclimatic variables selected for each species are shown in Table 4.4. AUC test and training values for all the species are above 0.9 (Table 4.5), this indicates that the potential distribution of all the species fits well with the data. For the majority of the species, precipitation seems to be the highest contributing factor towards their predicted distribution, although temperature also governs the distribution of some of the primate species in northeast India (Table 4.4). By looking at the SDM outputs from Maxent analysis we can see that, except for *Hoolock hoolock*, for most of the species their potential distribution increased in the Pleistocene (~787 Ka) period as compared to what it was in the Pliocene (~3.3 Ma). Then finally in the current time period (1979-2013), we can see that the potential distribution of most of the species decreased as compared to their distribution during the Pleistocene period. There were exceptions to this result, the potential distribution of *T. phayrei* and *N*. *bengalensis* shows a slight increase as compared to their distribution during Pleistocene, and the distribution of *H*. *hoolock* doesn’t show any change between the current and the Pleistocene periods.

## Discussion

In this study, our aim was to correlate the past population size changes in individual species, to the past climate changes, specifically mid Pliocene and warm Pleistocene interglacial. We found that each species had its own response to the environmental changes. We will be discussing these genus-wise to provide the clear implications from the study.

*A*. Genus – *Macaca* *M. assamensis* and *M. thibetana* both belong to the *sinica* group of macaques, along with *M. sinica, M. radiata, M. munzala, M. leucogenys* and *M. sela*. In our MSMC2 plots, both *M. assamensis* and *M. thibetana* have similar trajectories till about 800,000 years ago. This reflects their genetic history, as they are phylogenetically closest species which have diverged as recently as 750,000 years ago (Roos et al., 2019). This is reflected in the plots, as we can observe the effective population sizes are identical till the mid-Pleistocene boundary and then both the species diverge, suggesting speciation in the common ancestor. The distribution models provide us with a geographic indication of this process. The most suitable habitat for both the species in Pliocene is the present-day northeast India. From there, by the Pleistocene the spread has taken opposite directions. *M. assamensis* have gone towards the west, making its way into the Himalayan foothills. *M. thibetana* has taken the route towards northeast with Tibet and southern China as its spread, along with suitability shown in Nepal too, overlapping *M. assamensis*. This becomes clearer as we come to present where the east-west distribution of *M. assamensis* and *M. thibetana* are distinct (Choudhury, 2022; Khanal et al., 2018). Since the common point to divergence is northeastern India, we suggest it must be the location where the speciation of both the species could have taken place, between 700,000 and 800,000 years ago. *M. arctoides* (Stump tailed macaque) is another unique species which has been variously classified as either the member of the *fascicularis* group, or *mulatta* group or as a sole member of its own *arctoides* group. Its evolution has also been an issue of intense debate in the community. *M. arctoides* diverged before the divergence of *M. assamensis* and *M. thibetana;* and later at 3.43 million years ago from *mulatta* group (Li et al., 2009; Roos et al., 2019). We see a peak for effective population size in our *M. arctoides* MSMC plot at about 1.3-1.4 million years ago and then there is steady decline regardless of climatic conditions. Observing the distribution models, we can see that at all the three time points the population is confined to the mainland southeast Asia. Other relatives of *M. arctoides* are present at both the north and south of this distribution, like *M. cyclopis* in Taiwan and *M. fuscata* in Japan. It has also been strongly suggested that *M. arctoides* is a result of hybridization between a proto-*arctoides* and female *M. mulatta* (Fan et al., 2018). Taking into account all this evidence, we speculate that a proto- *arctoides* ancestor could have evolved in the Indochina region and then various populations would have proceeded to different regions, decreasing the effective population size at about 1.3-1.4 million years ago. Following this, a population might have moved northward and hybridised with *M. mulatta* to give rise to modern *M. arctoides*. Another species we considered was *M. leonina* (Northern Pig tailed macaque). This is a sister species of *M. nemestrina*, the southern pig-tailed macaque, which also lends its name to the *nemestrina* group, which includes all the Sulawesi macaque species and *M. silenus* (lion-tailed macaque) in peninsular India. This is the oldest diverged group of macaques, after their initial split from *M. sylvanus* (Barbary macaque) (Roos et al., 2019). As with *M. arctoides*, the distribution models suggest that *M. leonina* also had suitable climates in southeast Asia, during Pliocene and Pleistocene, and later it must have expanded its range towards northeast India. In the MSMC plots, there are two drastic population decline, one at 2 million years ago and another at 350,000 years ago. We expect that there could have been a speciation event at 2 mya point, as this is supported by the literature. This could be the point when both southern and northern pig-tailed macaque species split somewhere in Indochina, with proto-*leonina* continuing its march to the north. Another splitting event, the most recent one, could have taken place during Pleistocene glaciations, which would have given the modern species of *M. leonina* and *M. silenus. M. silenus* could have formed from a population of *M. leonina* which got separated during glaciations and went towards peninsular India in the search of more conducive environments. After the Pleistocene, the contact between both the population was lost due to lack of continuous forests, following the allopatric speciation of *M. silenus* (Ram et al., 2015; Singh et al., 2002). A similar study of *M. silenus* would be immensely beneficial to support or refute these conjectures.
*B*. Genus – *Trachypithecus* *Trachypithecus* are leaf monkeys which are related to more common hanuman langurs, which form the genus *Semnopithecus. Trachypithecus* consists of about 16 species, out of which three are found in northeast India: *Trachypithecus pileatus* (capped langur)*, T. geei* (golden langur) and *T. phayrei* (Phayre’s leaf monkey) (He et al., 2012; Roos et al., 2020; Wang et al., 2012). We were able to get samples and location data for all the three species and analysed demography and historical distribution for them. Though there is a lack of a conclusive phylogeny, mitochondrial data has shown that the ancestor of *T. pileatus* and *T. geei* was the first one to split in the *Trachypithecus* genus at about 4.5 million years ago (Roos et al., 2020). The divergence between *T. pileatus* and *T. geei* is much recent, at about 500,000 to 800,000 years ago. We find a peak in the effective population size of *T. pileatus* exactly at the 3.3 million years, showing that this species preferred warmer climate of mid-Pliocene. There is a drastic decline as the Pleistocene glaciations start, only to rise and then again declining at 787 kya mark. Here, we expect the divergence of *T. geei* from *T. phayrei*. A corroboration also comes from the demography of *T. geei*. Here the peak is at the 787 kya point, where the *T. pileatus* population goes in decline, suggesting that some population might have speciated to *T. geei* during this period. In the distribution maps, in both Pliocene and Pleistocene, the population in all the scenarios seems to always concentrated in northeast India and Bangladesh for *T. phayrei*. There is spread towards Myanmar in mid-Pleistocene in which can indicate distribution of the common ancestor of *T. pileatus* and *T. geei*. From there, some population of (or related) to *T. pileatus* could have seek refugia in the hills of present-day Assam-Bhutan border and then have evolved into *T. geei*. Hybridization between Hanuman langurs (*Semnopithecus* spp.) and *T. pileatus* has also been proposed as a reason for the speciation of *T. geei* as the present distribution of the species falls into hybridization zone of both Hanuman langurs and capped langur (Arekar et al., 2021). *T. phayrei* is a member of a different group, *obscurus*, within the lutungs which have recently diverged into four species. The split of *T. phayrei* from its ancestral group has happened at 1.4 million years ago according to the mitochondrial data (Roos et al., 2020). In our demographic estimates, we find the peak population at 1-million-year mark, which then steadily declines after mid-Pleistocene. After the split, *T. phayrei* looks to have survived the ice ages with no further divergence in new species. If we look at historical distribution maps, it is evident that *T. phayrei* also has had a constant ancestral distribution only in northeast India and Bangladesh, and then it has slowly distributed towards southeast Asia. Seeing the pattern of the three *Trachypithecus* species in northeast India, we can hypothesize that the region can be a ‘centre of origin’ for the ancestral *Trachypithecus* species and might also be for the proto-*Trachypithecus-Semnopithecus* species. This idea has also been suggested by some other authors based on molecular work (Karanth, 2010; Osterholz et al., 2008; Roos et al., 2011).
C. Genus – *Nycticebus* *Nycticebus* is the genus of slow lorises, which contain eight species, distributed all over the southeast Asia. *Nycticebus bengalensis* is the single species found in northeast India; it is also the one with western most distribution in the genus. *Nycticebus* are the oldest genus among lorisids and their ancestor can be traced back till Miocene. The basal lineage of *N. pygmaeus* seems to be diverged at 11 million years ago (Chen et al., 2006; Munds et al., 2018). According to the same mitochondrial analysis, *N. bengalensis* is one of the most recent one, with the last split from *N. javanicus* in Pleistocene. The lack of data for such old conditions prevents us from commenting anything about the evolution of the whole genus. Our distribution models indicate the probability of the presence of *N. bengalensis* in northeast India at both the time points of Pliocene and Pleistocene. Observing the demographic history, we can say that *N. bengalensis* thrived in mid Pleistocene, and then suddenly declined as the glaciations proceeded. We expect this can be the time when the last split between *N. bengalensis* and *N. javanicus* could have happened, though this speculation has to be taken with a grain of salt. Present distributions of *N. bengalensis* and *N. javanicus* are not continuous, as the *N. coucang* lies in the middle of them. This disjoint distribution makes it difficult to guess any historical biogeographic scenario, especially when location and genetic data of most species in the genus are scarce. We can just strongly suggest that *N. bengalensis* or its ancestor had a suitable habitat in northeast India since at least Pliocene, and then could have gone under population bottleneck and speciation during the repeated glacial periods of Pleistocene.
D. Genus – *Hoolock* Gibbons are the small apes, among which *Hoolock* is the genus found in India, with *Hoolock hoolock* (Western hoolock gibbon) as its sole representative. Gibbon diverged from great apes at about 16-17 million years ago and then the radiation of gibbon family into four genera began at ~6 million years ago (Carbone et al., 2014; Trivedi et al., 2021). Hylobatidae phylogeny is still a matter of debate and is pending to be resolved even after extensive research using various datasets and techniques. In our demographic and distribution history, the conclusions seem to be marred by these confusions about gibbon evolution. The effective population size of *H. hoolock* has a small plateaued peak at mid-Pliocene and then a steep rise in population after mid-Pleistocene followed by a drastic decline. Therefore, the warmer period of Pliocene has suited the gibbons, but increased oscillations of Pleistocene in comparatively recent history seem to be averse to their preference of habitat. Observing the distribution models, we see similar extent in all the three periods. There is a small distribution in the areas of southern Assam, Meghalaya and Bangladesh. Another place of distribution is in the upper Assam. It is shown as disjoint by the model. As the time has gone by, there is a slight decrease in the southern distribution and a slight increase in the northern one. This change in the distribution cannot be termed as significant enough to give strong conclusions about the *H. hoolock* evolutionary history.

In this study, we have assimilated the genomic and presence data of nine species of primates in the northeast India. Our aim was to understand the speciation in these species with details on temporal and spatial scale, to decipher their origin and their trajectory in or out from northeast India. We found that there are various timescales and routes of ingress or egress depending on the genus and species. In the genus *Macaca*, *M. assamensis* and *M. thibetana*, their ancestor seems to be in the upper northeast region and they diverged from there to present habitats in different directions; *M. arctoides* can be the result of hybridization between proto-*arctoides* and rhesus macaque, which could have happened in mainland Indochina, after proto-*arctoides* started marching towards north from coastal regions in southeast Asia; *M. arctoides* could have diverged from the populations that came from southeast Asia and got isolated during the glacial period. One of these populations could have continued towards peninsular India to evolve into *M. silenus*. In *Trachypithecus*, the origin of all the species seems to be from the northeast India. *T. Pileatus* was the first to diverge and retains its distribution in the region since its inception. During Pleistocene, one of the isolated populations evolved into *T. geei*. Whether this was due to an introgression by a *Semnopithecus* species is still an open question. *T. phayrei* also doesn’t seem to change much, at least in the northeast region and has retained its distribution. *Nycticebus bengalensis* is the oldest species and seems to have prospered well in both Pliocene and Pleistocene with not much alteration in population sizes during these periods. For *Hoolock hoolock*, our data does not allow us to provide any robust conclusions about its evolutionary history.

These findings suggests that northeast India provides enough opportunities for species evolution due to varied resource availability. It remains an immensely intriguing study system where the confluence of species has brought about a peculiar blend of communities. This study is a testimony to this and we strongly emphasize the need for an extensive future research programme to understand primates and other taxa, which will consecutively will assist in better conservation of the region and its biodiversity.

**Figure 1.**
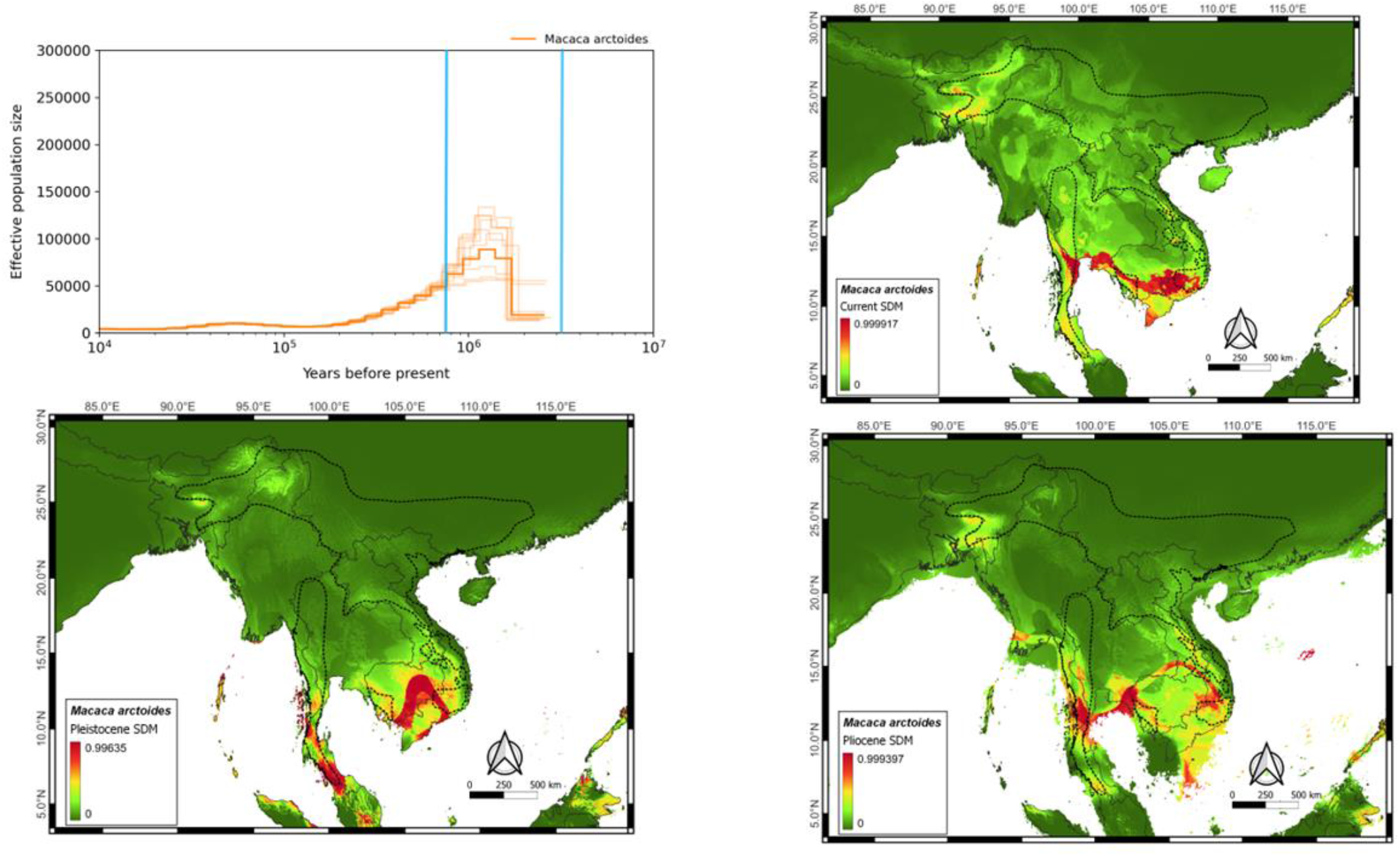
Macaca arctoides. Clockwise, the figure shows effective population size from about 2.5 million years ago to the present. Next one is the modelled present distribution showing majority of population in southeaster Indochina. Then is the modelled distribution in Pliocene (3.3 million years ago) showing disparate populations in Indochina and last one is Pleistocene (787,000 years ago) demonstrating not much change in distribution from Pliocene.

**Figure 2.**
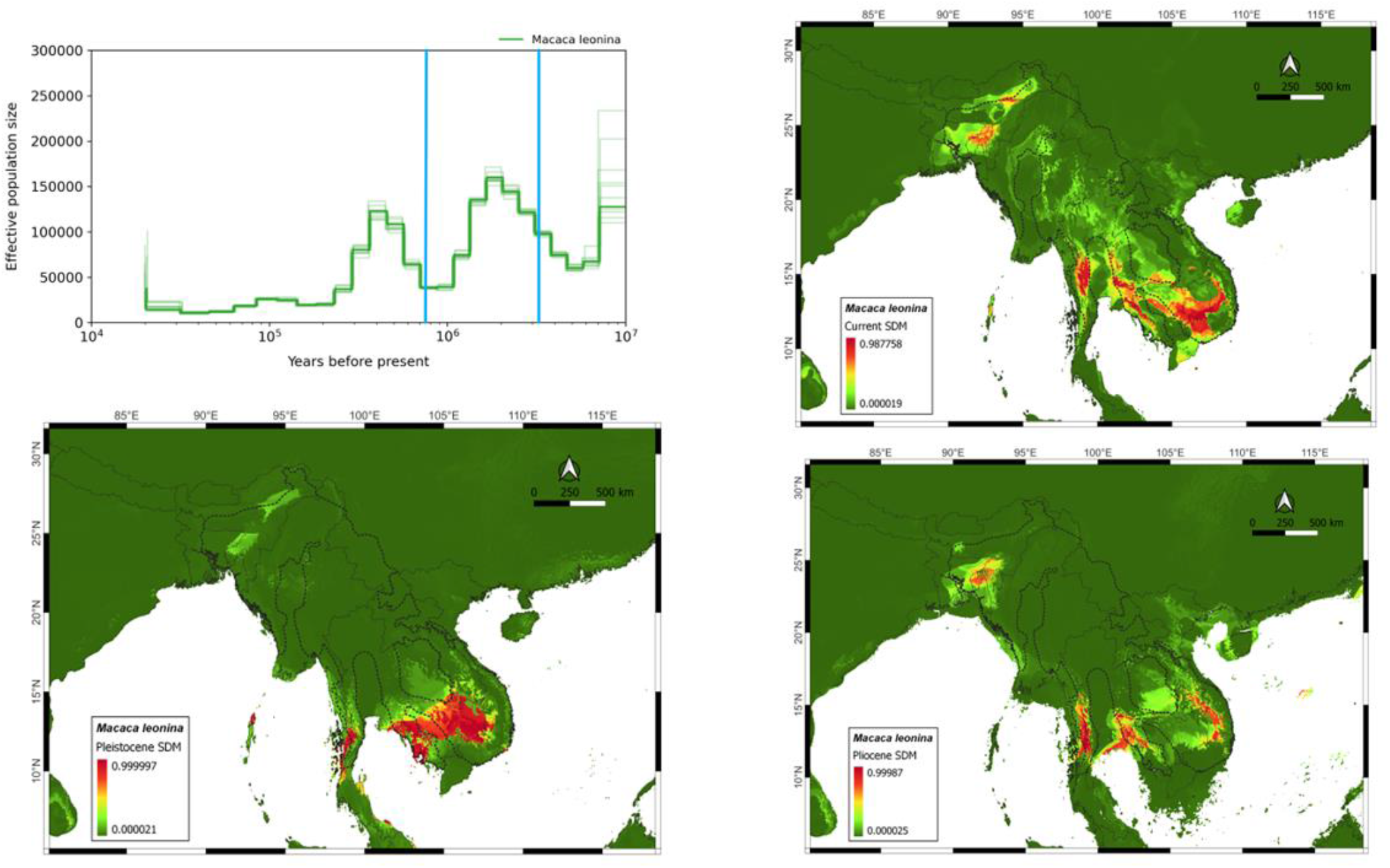
Macaca leonina. Clockwise, the figure shows effective population size from about 10 million years ago to the present, though before 8 million years is not significant. Next one is the modelled present distribution showing majority of population in most of mainland southeast Asia. Modelled distribution in Pliocene (3.3 million years ago) showing disparate populations in Indochina and last one is Pleistocene (787,000 years ago) has only one large population, with very less probability in northeast India.

**Figure 3.**
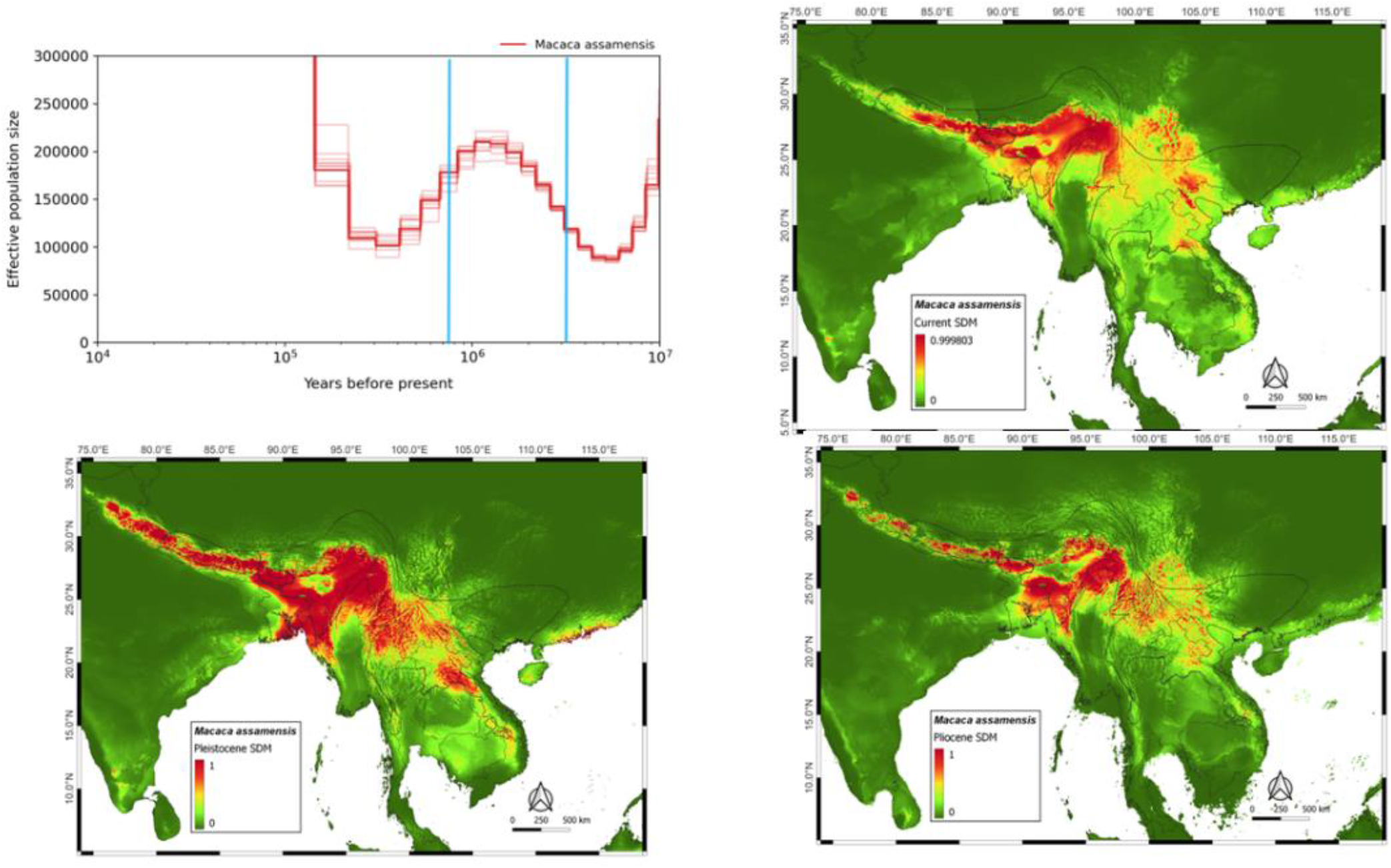
Macaca assamensis. Clockwise, the figure shows effective population size from about 10 million years ago to the present. The Modelled present distribution shows the whole expansion from Nepal to Vietnam. Modelled distribution in Pliocene (3.3 million years ago) shows a concentrated population in Northeast India and last one is Pleistocene (787,000 years ago) this northeastern population has spread, *en route* to achieving the present distribution.

**Figure 4.**
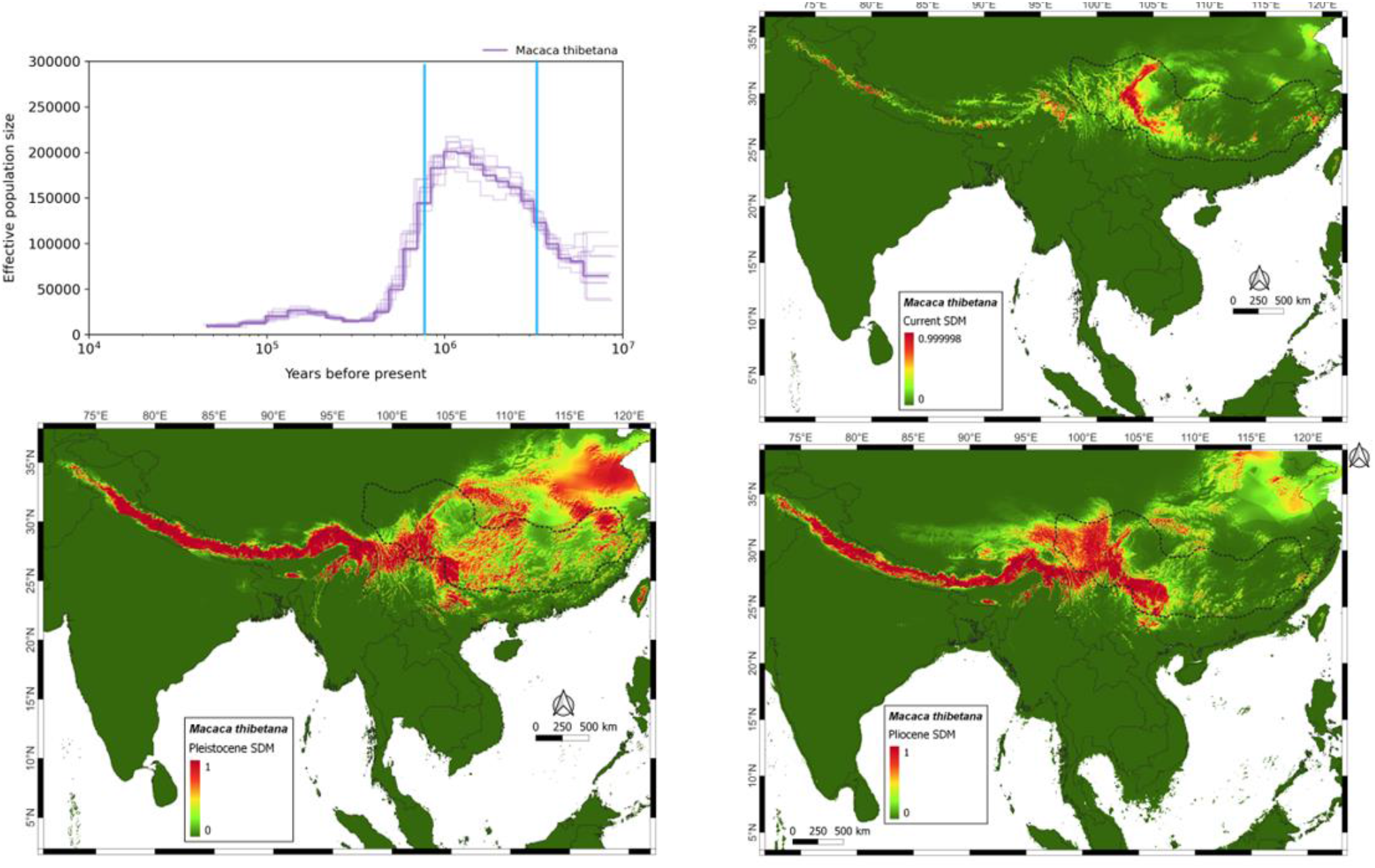
Macaca thibetana. Clockwise, the figure shows effective population size from about 8 million years ago to the present. The Modelled present distribution shows the present population is concentrated in southern Tibet. Modelled distribution in Pliocene (3.3 million years ago) shows a concentrated population in Northeast India and adjoining parts of Tibet. Pleistocene (787,000 years ago) is showing that this Pliocene population has spread towards the east.

**Figure 5.**
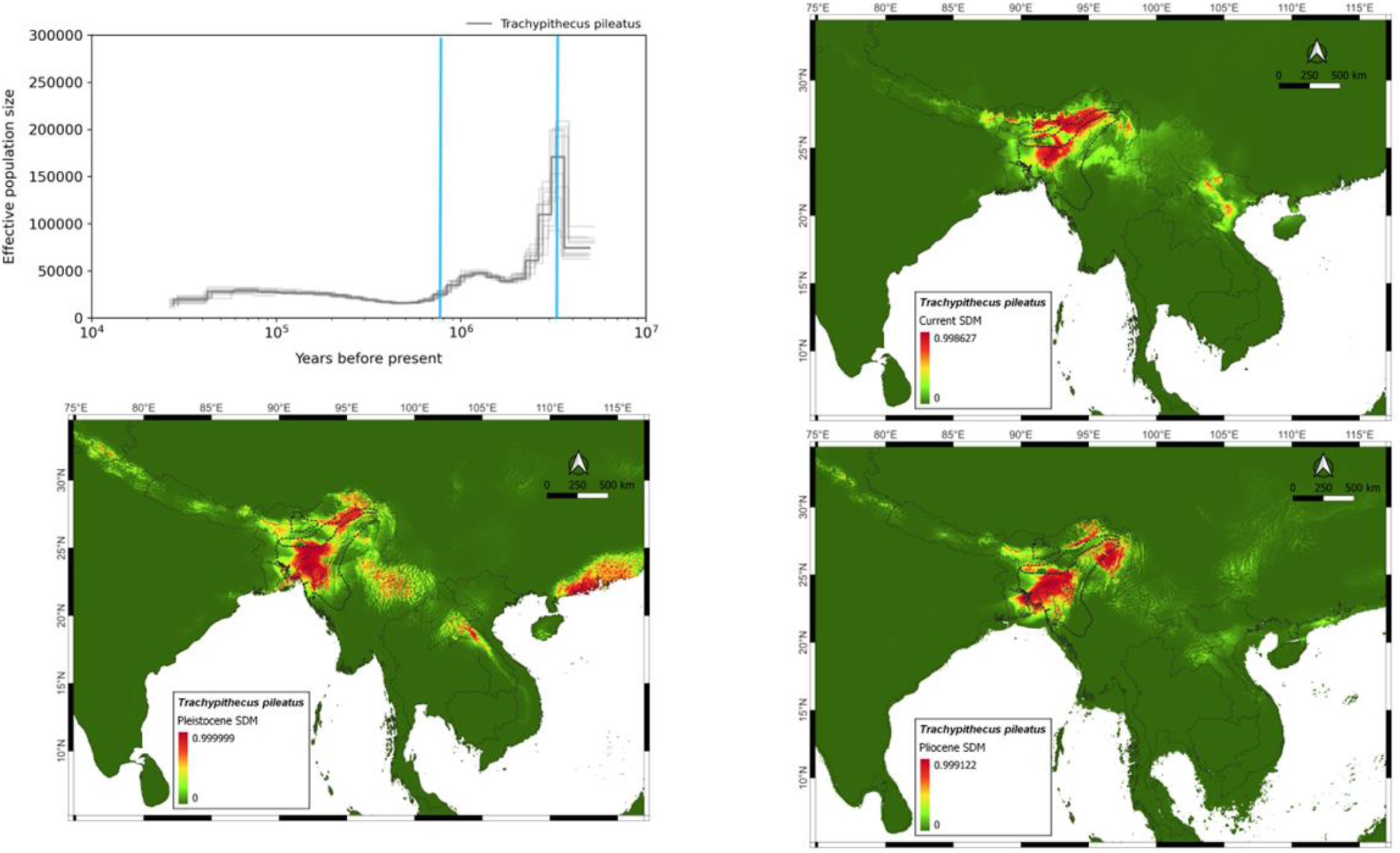
Trachypithecus pileatus. Clockwise, the figure shows effective population size from about 4 million years ago to the present. The Modelled present distribution shows the present population is concentrated in northeast India. Modelled distribution in Pliocene (3.3 million years ago) and Pleistocene (787,000 years ago) shows that the population has always been there in the region with very less variation with time.

**Figure 6.**
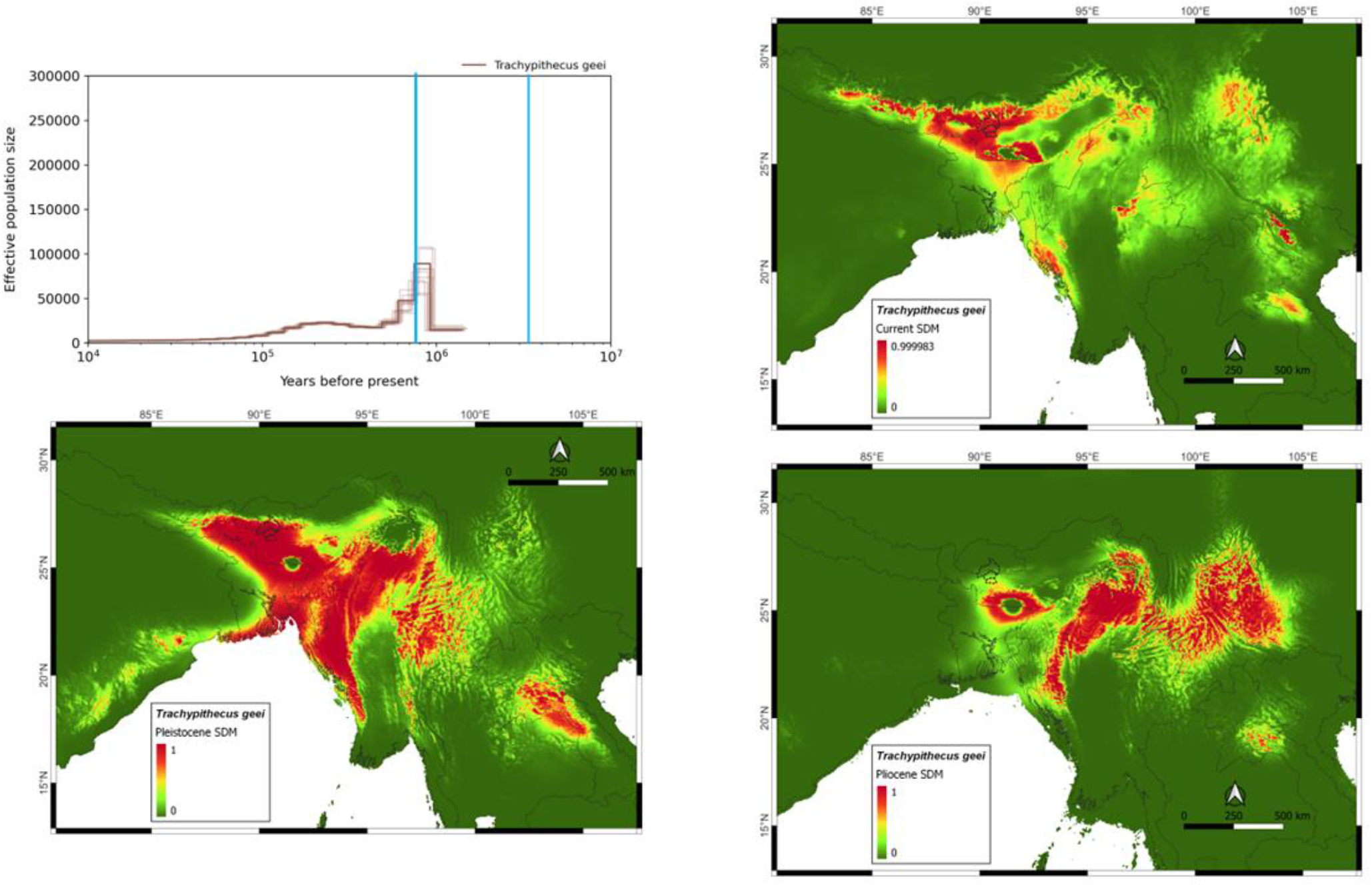
Trachypithecus geewe. Clockwise, the figure shows effective population size from about 1.5 million years ago to the present. The Modelled present distribution shows the present population is concentrated in northeast India. Modelled distribution in Pliocene (3.3 million years ago) shows some population in Ouranmar and Southern China. Pleistocene (787,000 years ago) shows that the Pliocene population has spread towards the northeast India.

**Figure 7.**
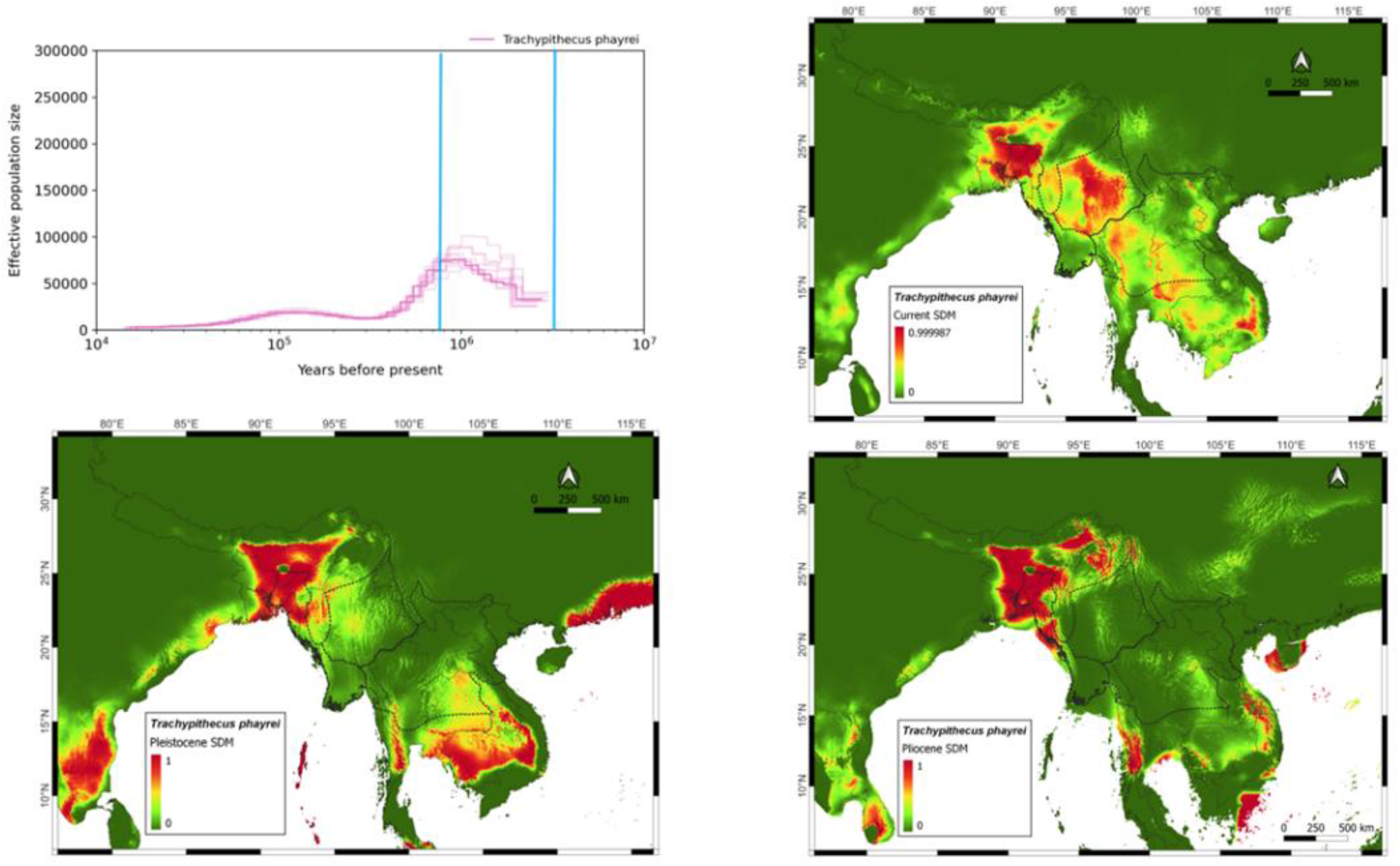
Trachypithecus phayrewe. Clockwise, the figure shows effective population size from about 3 million years ago to the present. The Modelled present distribution shows the present distribution is all over mainland southeast Asia. Modelled distribution in Pliocene (3.3 million years ago) shows that was concentrated to Northeast India. Pleistocene (787,000 years ago) shows that the Pliocene population had started spreading towards southeast Asia.

**Figure 8.**
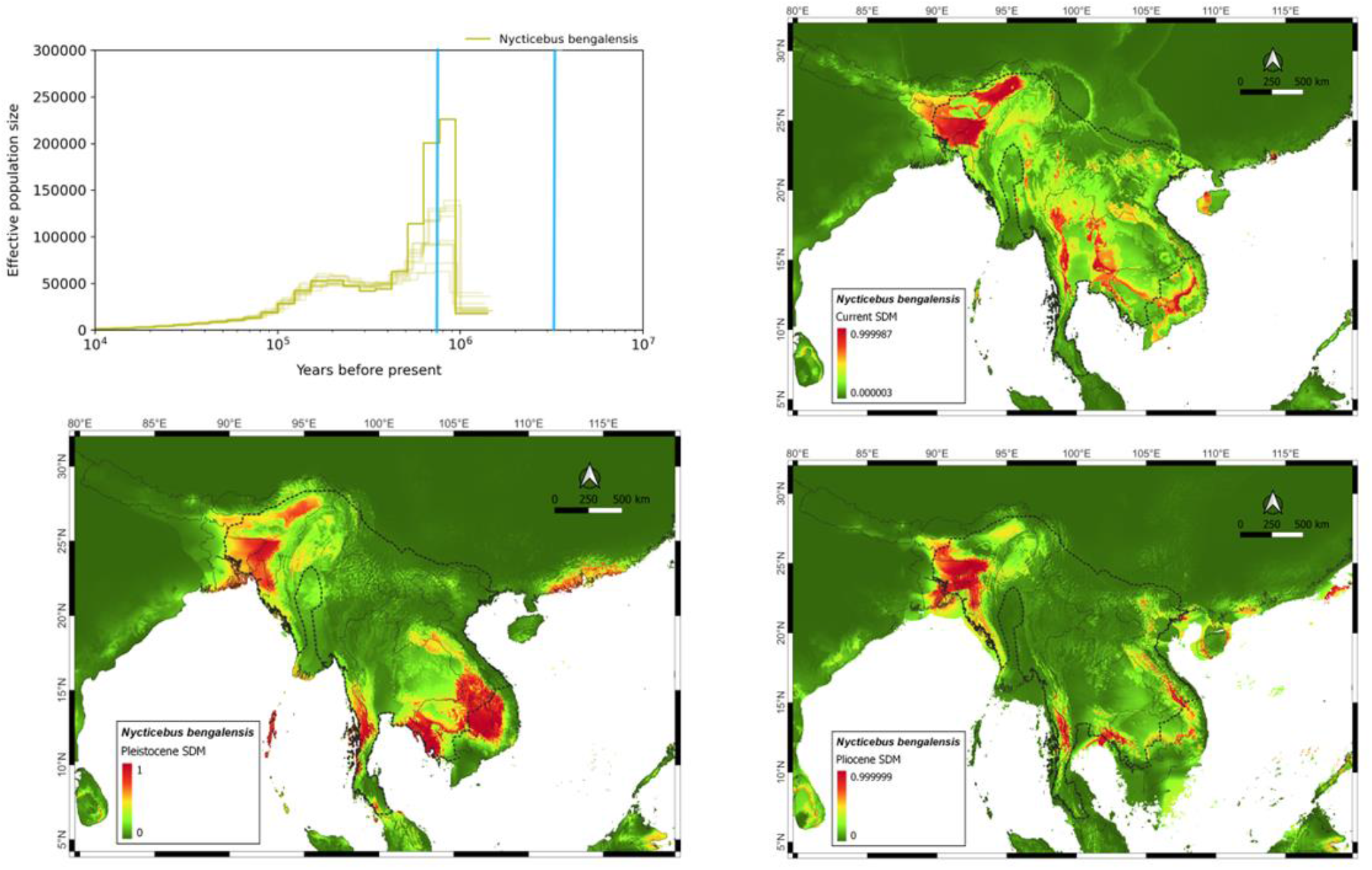
Nycticebus bengalensis. Clockwise, the figure shows effective population size from about 1 million years ago to the present. The Modelled present distribution shows the present distribution is all over mainland southeast Asia. Modelled distribution in Pliocene (3.3 million years ago) shows that was concentrated to Northeast India. Pleistocene (787,000 years ago) has the evidence of a small population in southeast Indochina.

**Figure 9.**
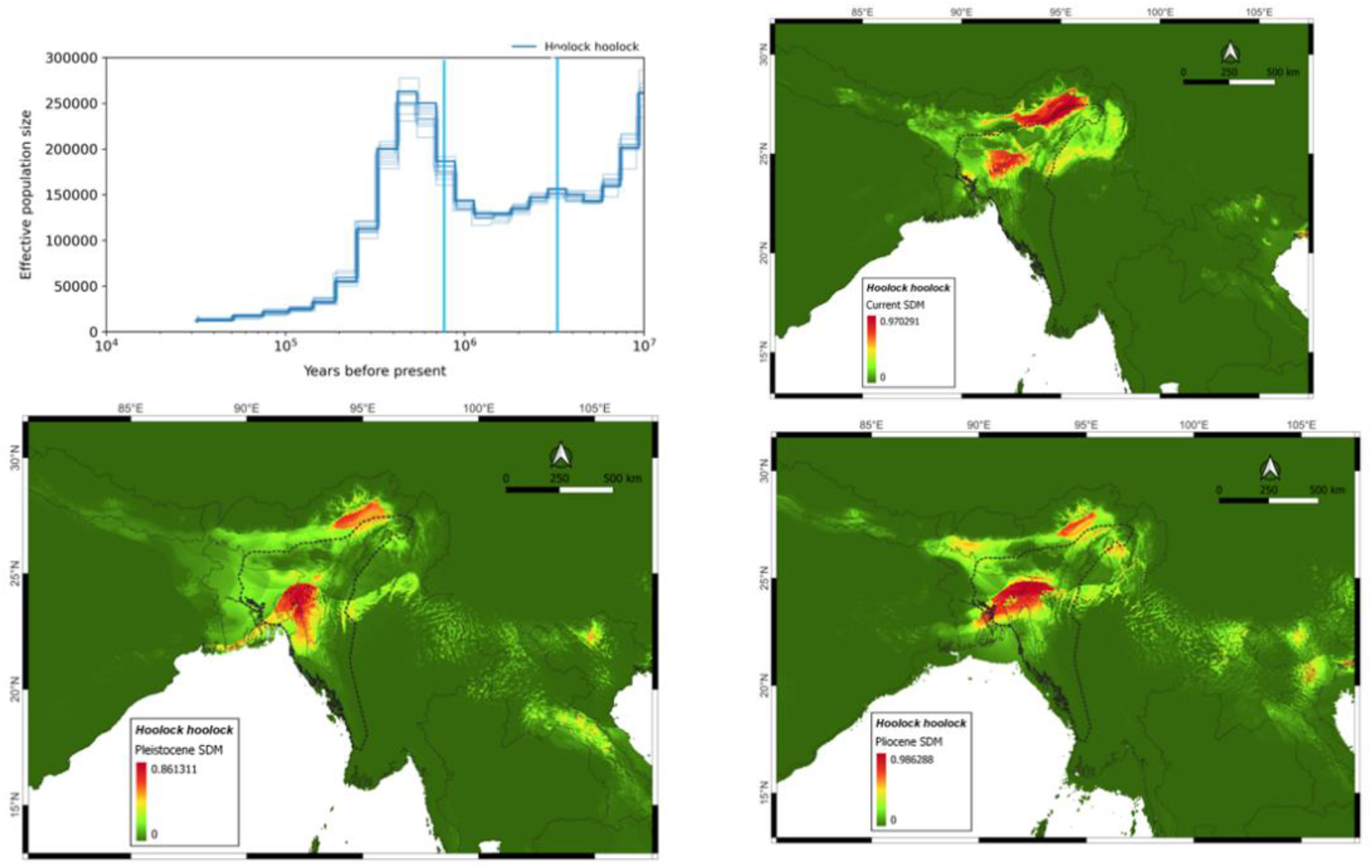
Hoolock hoolock. Clockwise, the figure shows effective population size from about 1 million years ago to the present. The Modelled present distribution shows the present distribution isis concentrated in northeast India. Modelled distribution in Pliocene (3.3 million years ago) and Pleistocene (787,000 years ago) both are showing similar distributions.

**Figure 10.**
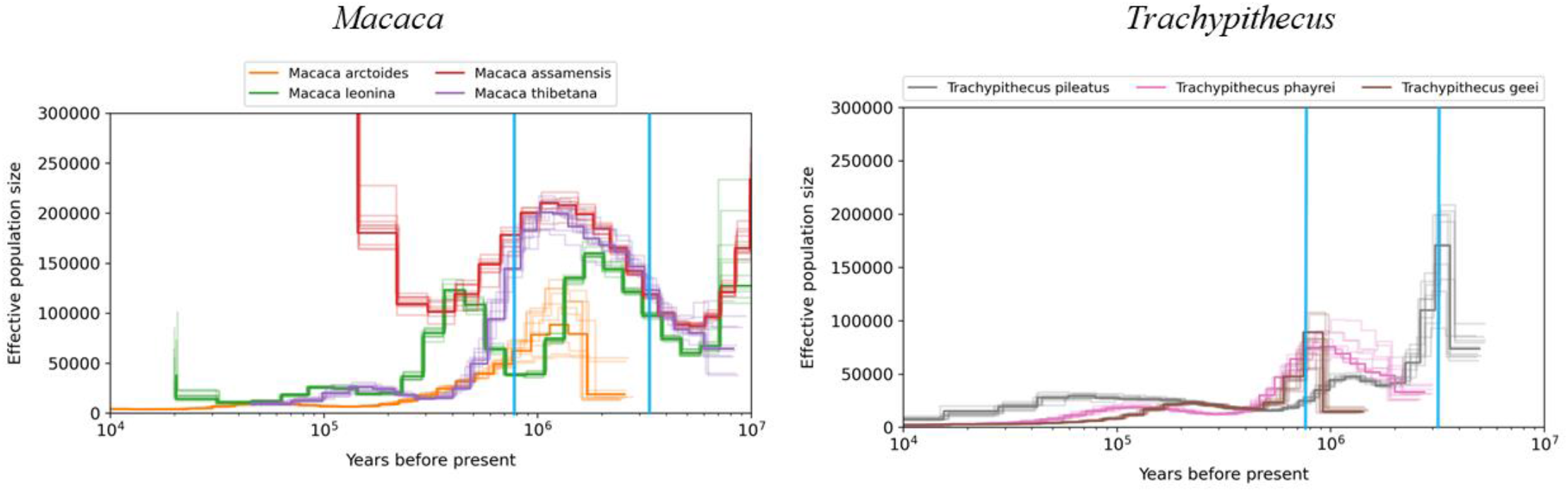
Assimilated MSMC graphs for genera *Macaca* and *Trachypithecus* for comparison.

**Table 1:** Number of individuals for each species

| S. No. | Species | Number of individuals | Reference species |
| --- | --- | --- | --- |
| 1 | *Macaca arctoides* | 4 | *Macaca mulatta* |
| 2 | *M. assamensis* | 1 | *M. mulatta* |
| 3 | *M. thibetana* | 1 | *M. mulatta* |
| 4 | *M. leonina* | 5 | *M. mulatta* |
| 5 | *Trachypithecus phayrei* | 1 | *Trachypithecus phayrei* |
| 6 | *T. pileatus* | 1 | *T. phayrei* |
| 7 | *T. geei* | 6 | *T. phayrei* |
| 8 | *Hoolock hoolock* | 11 | *Nomascus leucogenys* |
| 9 | *Nycticebus bengalensis* | 4 | *Nycticebus bengalensis* |

**Table 2:**
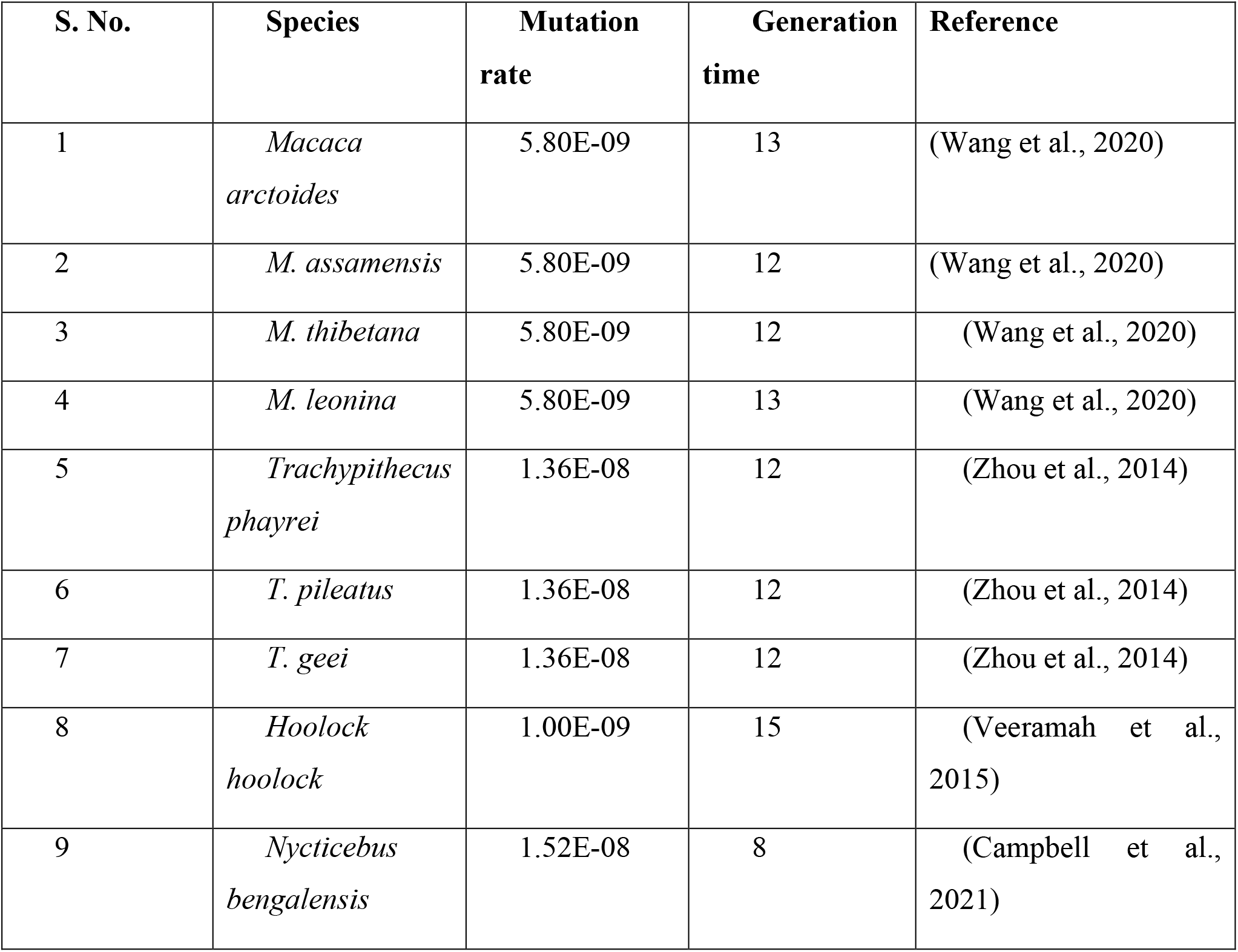
Mutation rate and generation time used by Ne estimation

| S. No. | Species | Mutation rate | Generation time | Reference |
| --- | --- | --- | --- | --- |
| 1 | *Macaca arctoides* | 5.80E-09 | 13 | (Wang et al., 2020) |
| 2 | *M. assamensis* | 5.80E-09 | 12 | (Wang et al., 2020) |
| 3 | *M. thibetana* | 5.80E-09 | 12 | (Wang et al., 2020) |
| 4 | *M. leonina* | 5.80E-09 | 13 | (Wang et al., 2020) |
| 5 | *Trachypithecus phayrei* | 1.36E-08 | 12 | (Zhou et al., 2014) |
| 6 | *T. pileatus* | 1.36E-08 | 12 | (Zhou et al., 2014) |
| 7 | *T. geei* | 1.36E-08 | 12 | (Zhou et al., 2014) |
| 8 | *Hoolock hoolock* | 1.00E-09 | 15 | (Veeramah et al., 2015) |
| 9 | *Nycticebus bengalensis* | 1.52E-08 | 8 | (Campbell et al., 2021) |

**Table 3:** Feature class type and RM values selected for the 9 species after the model selection analysis. FC = Feature class; RM = Regularisation multiplier

| **Species** | **FC** | **RM** |
| --- | --- | --- |
| *Hoolock hoolock* | LQPT | 1 |
| *Macaca arctoides* | LQPT | 1 |
| *Macaca assamensis* | QPT | 2 |
| *Macaca leonina* | T | 1 |
| *Macaca thibetana* | QPT | 1 |
| *Nycticebus bengalensis* | QPT | 1 |
| *Trachypithecus geei* | LQ | 0.5 |
| *Trachypithecus phayrei* | LQP | 0.5 |
| *Trachypithecus pileatus* | QPTH | 1 |

**Table 4:** Bioclimatic variables selected for each of the 9 species for Palaeodistribution modelling in Maxent.

| *H.*<br>*hoolock* | *M.*<br>*arctoides* | *M.*<br>*assamensis* | *M.*<br>*leonina* | *M.*<br>*thibetana* | *N.*<br>*bengalensis* | *T.*<br>*geei* | *T.*<br>*phayrei* | *T.*<br>*pileatus* |
| --- | --- | --- | --- | --- | --- | --- | --- | --- |
| bio1 | bio1 | bio1 | bio1 | bio4 | bio1 | bio1 | bio1 | bio1 |
| bio4 | bio4 | bio4 | bio4 | bio11 | bio4 | bio4 | bio4 | bio4 |
| bio12 | bio12 | bio13 | bio12 | bio12 | bio12 | bio13 | bio12 | bio13 |
| bio15 | bio14 | bio15 | bio14 | bio14 | bio14 | bio15 | bio14 | bio14 |
| bio17 | bio15 | bio17 | bio15 | bio15 | bio15 | bio17 | bio15 | bio15 |
| bio18 | bio18 | bio18 | bio18 | bio18 | bio18 | bio18 | bio18 | bio18 |

*H* = *Hoolock*, *M* = *Macaca*, *N* = *Nycticebus*, *T* = *Trachypithecus*

**Table 5:** AUC test, AUC training and top three bioclimatic variables showing the highest contribution to the model.

| Species | AUC<br>Test | AUC<br>Training | Variable | Percent<br>contribution | Permutation<br>importance |
| --- | --- | --- | --- | --- | --- |
| *Hoolock hoolock* | 0.99 | 0.993 | bio18.tif | 64.8 | 1.7 |
|  |  |  | bio4.tif | 13.5 | 21.1 |
|  |  |  | bio1.tif | 6.9 | 57.1 |
| *Macaca arctoides* | 0.951 | 0.975 | bio4.tif | 65 | 76.3 |
|  |  |  | bio14.tif | 16.7 | 8.2 |
|  |  |  | bio15.tif | 9.5 | 4.5 |
| *Macaca assamensis* | 0.954 | 0.96 | bio4.tif | 42.9 | 44.6 |
|  |  |  | bio18.tif | 22.6 | 2.2 |
|  |  |  | bio17.tif | 14.2 | 26.6 |
| *Macaca leonina* | 0.98 | 0.988 | bio15.tif | 37.1 | 54.5 |
|  |  |  | bio4.tif | 34 | 15.4 |
|  |  |  | bio12.tif | 10.1 | 8.6 |
| *Macaca thibetana* | 0.988 | 0.992 | bio11.tif | 48 | 82.1 |
|  |  |  | bio4.tif | 27.6 | 9.3 |
|  |  |  | bio18.tif | 14.5 | 7.6 |
| *Nycticebus bengalensis* | 0.911 | 0.98 | bio15.tif | 34.7 | 36.7 |
|  |  |  | bio18.tif | 23.9 | 6.5 |
|  |  |  | bio4.tif | 22.2 | 33.4 |
| *Trachypithecus geei* | 0.989 | 0.99 | bio18.tif | 48.2 | 0.9 |
|  |  |  | bio4.tif | 21.4 | 41.5 |
|  |  |  | bio17.tif | 11.8 | 37.8 |
| *Trachypithecus phayrei* | 0.958 | 0.962 | bio4.tif | 35.1 | 19.5 |
|  |  |  | bio14.tif | 20.3 | 43.2 |
|  |  |  | bio15.tif | 20.3 | 21.6 |
| *Trachypithecus pileatus* | 0.987 | 0.992 | bio18.tif | 55.4 | 0.7 |
|  |  |  | bio4.tif | 28.6 | 29.2 |
|  |  |  | bio15.tif | 8.3 | 23.8 |

## References

Aiello-Lammens, M. E., Boria, R. A., Radosavljevic, A., Vilela, B., & Anderson, R. P. (2015). spThin: an R package for spatial thinning of species occurrence records for use in ecological niche models. Ecography, 38(5), 541–545.

Arekar, K., Parigi, A., & Karanth, K. P. (2021). Understanding the convoluted evolutionary history of the capped-golden langur lineage (Cercopithecidae: Colobinae)†. Journal of Genetics, 100(2). https://doi.org/10.1007/s12041-021-01329-8

Barnosky, A. D., & Kraatz, B. P. (2007). The role of climatic change in the evolution of mammals. BioScience, 57(6), 523–532. https://doi.org/10.1641/B570615

Beyer, R. M., & Manica, A. (2020). Historical and projected future range sizes of the world’s mammals, birds, and amphibians. Nature Communications, 11(1), 1–8.

Briggs, J. C. (1987). Biogeography and plate tectonics. Elsevier.

Brown, J. L., Hill, D. J., Dolan, A. M., Carnaval, A. C., & Haywood, A. M. (2018). Paleoclim, high spatial resolution paleoclimate surfaces for global land areas. Scientific Data, 5, 1–9. https://doi.org/10.1038/sdata.2018.254

Campbell, C. R., Tiley, G. P., Poelstra, J. W., Hunnicutt, K. E., Larsen, P. A., Lee, H.-J., Thorne, J. L., Dos Reis, M., & Yoder, A. D. (2021). Pedigree-based and phylogenetic methods support surprising patterns of mutation rate and spectrum in the gray mouse lemur. Heredity, 127(2), 233–244.

Carbone, L., Alan Harris, R., Gnerre, S., Veeramah, K. R., Lorente-Galdos, B., Huddleston, J., Meyer, T. J., Herrero, J., Roos, C., Aken, B., Anaclerio, F., Archidiacono, N., Baker, C., Barrell, D., Batzer, M. A., Beal, K., Blancher, A., Bohrson, C. L., Brameier, M., … Gibbs, R. A. (2014). Gibbon genome and the fast karyotype evolution of small apes. Nature, 513(7517), 195–201. https://doi.org/10.1038/nature13679

Chen, J. H., Pan, D., Groves, C., Wang, Y. X., Narushima, E., Fitch-Snyder, H., Crow, P., Thanh, V. N., Ryder, O., Zhang, H. W., Fu, Y. X., & Zhang, Y. P. (2006). Molecular phylogeny of Nycticebus inferred from mitochondrial genes. International Journal of Primatology, 27(4), 1187–1200. https://doi.org/10.1007/s10764-006-9032-5

Chen, Z., Li, H., Zhai, X., Zhu, Y., He, Y., Wang, Q., Li, Z., Jiang, J., Xiong, R., & Chen, X. (2020). Phylogeography, speciation and demographic history: Contrasting evidence from mitochondrial and nuclear markers of the Odorrana graminea sensu lato (Anura, Ranidae) in China. Molecular Phylogenetics and Evolution, 144, 106701.

Choudhury, A. (2022). A Review of the Assamese Macaque Macaca assamensis Complex and Its Geographical Variation. Primate Conservation, 36, 203–231.

Crispo, E., DiBattista, J. D., Correa, C., Thibert-Plante, X., McKellar, A. E., Schwartz, A. K., Berner, D., De León, L. F., & Hendry, A. P. (2010). The evolution of phenotypic plasticity in response to anthropogenic disturbance. Evolutionary Ecology Research, 12(1), 47–66.

Davis, M. B., & Shaw, R. G. (2001). Range shifts and adaptive responses to Quaternary climate change. Science, 292(5517), 673–679.

Dolan, A. M., Haywood, A. M., Hunter, S. J., Tindall, J. C., Dowsett, H. J., Hill, D. J., & Pickering, S. J. (2015). Modelling the enigmatic late Pliocene glacial event—Marine Isotope Stage M2. Global and Planetary Change, 128, 47–60.

Evans, L. J., Goossens, B., Davies, A. B., Reynolds, G., & Asner, G. P. (2020). Natural and anthropogenic drivers of Bornean elephant movement strategies. Global Ecology and Conservation, 22, e00906.

Fan, Z., Zhou, A., Osada, N., Yu, J., Jiang, J., Li, P., Du, L., Niu, L., Deng, J., Xu, H., Xing, J., Yue, B., & Li, J. (2018). Ancient hybridization and admixture in macaques (genus Macaca) inferred from whole genome sequences. Molecular Phylogenetics and Evolution, 127(March), 376–386. https://doi.org/10.1016/j.ympev.2018.03.038

Galante, P. J., Alade, B., Muscarella, R., Jansa, S. A., Goodman, S. M., & Anderson, R. P. (2018). The challenge of modeling niches and distributions for data-poor species: a comprehensive approach to model complexity. Ecography, 41, 726–736.

Guisan, A., & Thuiller, W. (2005). Predicting species distribution: Offering more than simple habitat models. Ecology Letters, 8(9), 993–1009. https://doi.org/10.1111/j.1461-0248.2005.00792.x

Hao, T., Elith, J., Guillera-Arroita, G., & Lahoz-Monfort, J. J. (2019). A review of evidence about use and performance of species distribution modelling ensembles like BIOMOD. Diversity and Distributions, 25(5), 839–852. https://doi.org/10.1111/ddi.12892

Harvey, M. G., Singhal, S., & Rabosky, D. L. (2019). Beyond reproductive isolation: demographic controls on the speciation process. Annual Review of Ecology, Evolution, and Systematics, 50(1), 75–95.

Haywood, A. M., Dowsett, H. J., Valdes, P. J., Lunt, D. J., Francis, J. E., & Sellwood, B. W. (2009). Introduction. Pliocene climate, processes and problems. Philosophical Transactions of the Royal Society A: Mathematical, Physical and Engineering Sciences, 367(1886), 3–17. https://doi.org/10.1098/rsta.2008.0205

He, K., Hu, N. Q., Orkin, J. D., Nyein, D. T., Ma, C., Xiao, W., Fan, P. F., & Jiang, X. L. (2012). Molecular phylogeny and divergence time of Trachypithecus: with implications for the taxonomy of T. phayrei. Dong Wu Xue Yan Jiu = Zoological Research / “Dong Wu Xue Yan Jiu” Bian Ji Wei Yuan Hui Bian Ji, 33(E5–6), 104–110.https://doi.org/10.3724/sp.j.1141.2012.e05-06e104

Hewitt, G. (2000). The genetic legacy of the quaternary ice ages. Nature, 405(6789), 907–913. https://doi.org/10.1038/35016000

Hill, J. K., Griffiths, H. M., & Thomas, C. D. (2011). Climate change and evolutionary adaptations at species’ range margins. Annual Review of Entomology, 56, 143–159.

Hoffmann, A. A., & Sgró, C. M. (2011). Climate change and evolutionary adaptation. Nature, 470(7335), 479–485. https://doi.org/10.1038/nature09670

Hua, X., & Wiens, J. J. (2013). How does climate influence speciation? The American Naturalist, 182(1), 1–12.

Huey, R. B., Deutsch, C. A., Tewksbury, J. J., Vitt, L. J., Hertz, P. E., Álvarez Pérez, H. J., & Garland Jr, T. (2009). Why tropical forest lizards are vulnerable to climate warming. Proceedings of the Royal Society B: Biological Sciences, 276(1664), 1939–1948.

Jackson, S. T., & Overpeck, J. T. (2000). Responses of plant populations and communities to environmental changes of the late Quaternary. Paleobiology, 26(S4), 194–220.

Jaureguiberry, P., Titeux, N., Wiemers, M., Bowler, D. E., Coscieme, L., Golden, A. S., Guerra, C. A., Jacob, U., Takahashi, Y., Settele, J., & others. (2022). The direct drivers of recent global anthropogenic biodiversity loss. Science Advances, 8(45), eabm9982.

Karanth, K. P. (2010). Molecular systematics and conservation of the langurs and leaf monkeys of South Asia. Journal of Genetics, 89(4), 393–399. https://doi.org/10.1007/s12041-010-0057-3

Khanal, L., Chalise, M. K., He, K., Acharya, B. K., Kawamoto, Y., & Jiang, X. (2018). Mitochondrial DNA analyses and ecological niche modeling reveal post-LGM expansion of the Assam macaque (Macaca assamensis) in the foothills of Nepal Himalaya. American Journal of Primatology, 80(3), 1–13. https://doi.org/10.1002/ajp.22748

Lavergne, S., Mouquet, N., Thuiller, W., & Ronce, O. (2010). Biodiversity and climate change: Integrating evolutionary and ecological responses of species and communities. Annual Review of Ecology, Evolution, and Systematics, 41, 321–350. https://doi.org/10.1146/annurev-ecolsys-102209-144628

Lenoir, J., Hattab, T., & Pierre, G. (2017). Climatic microrefugia under anthropogenic climate change: implications for species redistribution. Ecography, 40(2), 253–266.

Li, J., Han, K., Xing, J., Kim, H.-S., Rogers, J., Ryder, O. A., Disotell, T., Yue, B., & Batzer, M. A. (2009). Phylogeny of the macaques (Cercopithecidae: Macaca) based on Alu elements. Gene, 448(2), 242–249.

Lieberman, B. S., & Eldredge, N. (1996). Trilobite biogeography in the Middle Devonian: geological processes and analytical methods. Paleobiology, 22(1), 66–79.

McKee, J. K. (2001). Faunal turnover rates and mammalian biodiversity of the late Pliocene and Pleistocene of eastern Africa. Paleobiology, 27(3), 500–511.

Midgley, G. F., Hannah, L., Millar, D., Rutherford, M. C., & Powrie, L. W. (2002). Assessing the vulnerability of species richness to anthropogenic climate change in a biodiversity hotspot. Global Ecology and Biogeography, 11(6), 445–451.

Mittelbach, G. G., Schemske, D. W., Cornell, H. V, Allen, A. P., Brown, J. M., Bush, M. B., Harrison, S. P., Hurlbert, A. H., Knowlton, N., Lessios, H. A., & others. (2007). Evolution and the latitudinal diversity gradient: speciation, extinction and biogeography. Ecology Letters, 10(4), 315–331.

Morales, N. S., Fernández, I. C., & Baca-González, V. (2017). MaxEnt’s parameter configuration and small samples: are we paying attention to recommendations? A systematic review. PeerJ, 5, e3093.

Munds, R. A., Titus, C. L., Eggert, L. S., & Blomquist, G. E. (2018). Using a multi-gene approach to infer the complicated phylogeny and evolutionary history of lorises (Order Primates: Family Lorisidae). Molecular Phylogenetics and Evolution, 127(October 2017), 556–567. https://doi.org/10.1016/j.ympev.2018.05.025

Neumann, L. K., Fuhlendorf, S. D., Davis, C. D., & Wilder, S. M. (2022). Climate alters the movement ecology of a non-migratory bird. Ecology and Evolution, 12(4), e8869.

Osterholz, M., Walter, L., & Roos, C. (2008). Phylogenetic position of the langur genera Semnopithecus and Trachypithecus among Asian colobines, and genus affiliations of their species groups. BMC Evolutionary Biology, 8(1), 1–12. https://doi.org/10.1186/1471-2148-8-58

Parmesan, C. (2006). Ecological and evolutionary responses to recent climate change. Annual Review of Ecology, Evolution, and Systematics, 37, 637–669. https://doi.org/10.1146/annurev.ecolsys.37.091305.110100

Parmesan, C., & Yohe, G. (2003). A globally coherent fingerprint of climate change. Nature, 421, 37–42.

Phillips, S. J., Anderson, R. P., & Schapire, R. E. (2006). Maximum entropy modeling of species geographic distributions. Ecological Modelling, 190(3–4), 231–259.

Poessel, S. A., Leitner, P., Inman, R. D., Esque, T. C., & Katzner, T. E. (2022). Demographic and environmental correlates of home ranges and long-distance movements of Mohave ground squirrels. Journal of Mammalogy.

Radosavljevic, A., & Anderson, R. P. (2014). Making better Maxent models of species distributions: Complexity, overfitting and evaluation. Journal of Biogeography, 41(4), 629–643. https://doi.org/10.1111/jbi.12227

Ram, M. S., Marne, M., Gaur, A., Kumara, H. N., Singh, M., Kumar, A., & Umapathy, G. (2015). Pre-historic and recent vicariance events shape genetic structure and diversity in endangered lion-tailed macaque in the Western Ghats: Implications for conservation. PLoS ONE, 10(11), 1–16.https://doi.org/10.1371/journal.pone.0142597

Roos, C., Helgen, K. M., Miguez, R. P., Thant, N. M. L., Lwin, N., Lin, A. K., Lin, A., Yi, K. M., Soe, P., Hein, Z. M., Myint, M. N. N., Ahmed, T., Chetry, D., Urh, M., Grace Veatch, E., Duncan, N., Kamminga, P., Chua, M. A. H., Yao, L., … Momberg, F. (2020). Mitogenomic phylogeny of the asian colobine genus trachypithecus with special focus on trachypithecus phayrei (Blyth, 1847) and description of a new species. Zoological Research, 41(6), 656–669. https://doi.org/10.24272/J.ISSN.2095-8137.2020.254

Roos, C., Kothe, M., Alba, D. M., Delson, E., & Zinner, D. (2019). The radiation of macaques out of Africa: Evidence from mitogenome divergence times and the fossil record. Journal of Human Evolution, 133, 114–132.

Roos, C., Zinner, D., Kubatko, L. S., Schwarz, C., Yang, M., Meyer, D., Nash, S. D., Xing, J., Batzer, M. A., Brameier, M., Leendertz, F. H., Ziegler, T., Perwitasari-Farajallah, D., Nadler, T., Walter, L., & Osterholz, M. (2011). Nuclear versus mitochondrial DNA: Evidence for hybridization in colobine monkeys. BMC Evolutionary Biology, 11(1). https://doi.org/10.1186/1471-2148-11-77

Salzmann, U., Haywood, A. M., Lunt, D. J., Valdes, P. J., & Hill, D. J. (2008). A new global biome reconstruction and data-model comparison for the Middle Pliocene. Global Ecology and Biogeography, 17(3), 432–447.

Shcheglovitova, M., & Anderson, R. P. (2013). Estimating optimal complexity for ecological niche models: A jackknife approach for species with small sample sizes. Ecological Modelling, 269, 9–17.

Singh, M., Singh, M., Kumar, M. A., Kumara, H. N., Sharma, A. K., & Kaumanns, W. (2002). Distribution, population structure, and conservation of lion-tailed macaques (Macaca silenus) in the Anaimalai Hills, Western Ghats, India. American Journal of Primatology: Official Journal of the American Society of Primatologists, 57(2), 91–102.

Smith, F. A., Elliott Smith, E. A., Villaseñor, A., Tomé, C. P., Lyons, S. K., & Newsome, S. D. (2022). Late Pleistocene megafauna extinction leads to missing pieces of ecological space in a North American mammal community. Proceedings of the National Academy of Sciences, 119(39), e2115015119.

Stillman, J. H. (2002). Causes and consequences of thermal tolerance limits in rocky intertidal porcelain crabs, genus Petrolisthes. Integrative and Comparative Biology, 42(4), 790–796.

Teixeira, H., Salmona, J., Arredondo, A., Mourato, B., Manzi, S., Rakotondravony, R., Mazet, O., Chikhi, L., Metzger, J., & Radespiel, U. (2021). Impact of model assumptions on demographic inferences: the case study of two sympatric mouse lemurs in northwestern Madagascar. BMC Ecology and Evolution, 21(1), 1–18. https://doi.org/10.1186/s12862-021-01929-z

Tierney, J. E., Poulsen, C. J., Montañez, I. P., Bhattacharya, T., Feng, R., Ford, H. L., Hönisch, B., Inglis, G. N., Petersen, S. V., Sagoo, N., Tabor, C. R., Thirumalai, K., Zhu, J., Burls, N. J., Foster, G. L., Goddéris, Y., Huber, B. T., Ivany, L. C., Turner, S. K., … Zhang, Y. G. (2020). Past climates inform our future. Science, 370(6517).https://doi.org/10.1126/science.aay3701

Trewick, S. (2017). Plate tectonics in biogeography. Int. Encycl. Geogr. People, Earth, Environ. Technol. (Eds Richardson D, Castree N, Goodchild M, Kobayashi A, Liu W, Marston R), 1–9.

Trivedi, M., Manu, S., Balakrishnan, S., Biswas, J., Asharaf, N. V. K., & Umapathy, G. (2021). Understanding the Phylogenetics of Indian Hoolock Gibbons: Hoolock hoolock and H. leuconedys. International Journal of Primatology, 42(3), 463–477.https://doi.org/10.1007/s10764-021-00212-8

Veeramah, K. R., Woerner, A. E., Johnstone, L., Gut, I., Gut, M., Marques-Bonet, T., Carbone, L., Wall, J. D., & Hammer, M. F. (2015). Examining phylogenetic relationships among gibbon genera using whole genome sequence data using an approximate bayesian computation approach. Genetics, 200(1), 295–308. https://doi.org/10.1534/genetics.115.174425

Wang, R. J., Thomas, G. W. C., Raveendran, M., Harris, R. A., Doddapaneni, H., Muzny, D. M., Capitanio, J. P., Radivojac, P., Rogers, J., & Hahn, M. W. (2020). Paternal age in rhesus macaques is positively associated with germline mutation accumulation but not with measures of offspring sociability. Genome Research, 30(6), 826–834.

Wang, X. P., Yu, L., Roos, C., Ting, N., Chen, C. P., Wang, J., & Zhang, Y. P. (2012). Phylogenetic relationships among the colobine monkeys revisited: New insights from analyses of complete mt genomes and 44 nuclear non-coding markers. PLoS ONE, 7(4), 1–12. https://doi.org/10.1371/journal.pone.0036274

Warren, D. L., Glor, R. E., & Turelli, M. (2010). ENMTools: a toolbox for comparative studies of environmental niche models. Ecography, 33(3), 607–611.

Williams, J. G., Zabel, R. W., Waples, R. S., Hutchings, J. A., & Connor, W. P. (2008). Potential for anthropogenic disturbances to influence evolutionary change in the life history of a threatened salmonid. Evolutionary Applications, 1(2), 271–285.

Zhou, X., Wang, B., Pan, Q., Zhang, J., Kumar, S., Sun, X., Liu, Z., Pan, H., Lin, Y., Liu, G., & others. (2014). Whole-genome sequencing of the snub-nosed monkey provides insights into folivory and evolutionary history. Nature Genetics, 46(12), 1303–1310.

